# Toward a Transferable Representation of Protein Dynamics for Functional Prediction and Mechanistic Discovery

**DOI:** 10.64898/2026.10.02.756378

**Authors:** Nahid Monowar, Lutfi Habiba

## Abstract

Protein function is determined not only by the static structure of molecules, but also by coordinated conformational fluctuations that occur across various spatial and temporal scales. Molecular dynamics simulations, elastic network models, principal component analysis, and related approaches offer useful descriptions of protein motion. However, representing heterogeneous trajectories in a compact, informative manner that is applicable to proteins with different sequences, structures, and historical evolution is not trivial.

Here we present the Topology Invariant Dynamic Graph Network (TIDGN), a deep representation learning approach that encodes protein conformational dynamics into a shared latent representation, termed the protein motion signature (Zp). TIDGN encodes each trajectory as a time-dependent residue-level graph and builds structural connectivity, dynamic cross-correlation, low-dimensional motion features, sequence context, and temporal aggregation into the design. Instead of learning a protein-specific collective variable, the approach learns a representation that retains functionally informative patterns of coordinated motion applicable to different proteins.

We evaluate the framework using 1,250 molecular-dynamics trajectories representing 1,120 distinct proteins that range in size from 50-600 residues, 142 CATH/SCOP families, 88 structural folds, and six broad functional classes. The set of trajectories spans durations of 100 ns to 10 µs, sampled at 100 ps intervals, representing approximately 12.5 million conformational frames in total. In the context of a sequence-disjoint evaluation using a 30% sequence-identity threshold, the approach achieves a mean AUROC of 0.8838 ± 0.0107 compared to 0.8174 for a dynamic graph neural-network baseline, 0.7257 for a dynamic descriptor, 0.7227 for a PCA descriptor, 0.6592 for a sequence baseline, and 0.6587 for a static-structure baseline. As the biological distance between training and evaluation proteins increases, transfer performance decreases with AUROC values of 0.92 for random splitting, 0.85 for sequence-disjoint evaluation, 0.78 for family-disjoint evaluation, and 0.71 for fold-disjoint evaluation.

Component ablations suggest that graph structure, dynamic cross-correlation, low-dimensional motion features, sequence information, and the combination of these factors contribute to predictive performance. Residue-level attribution, when applied to adenylate kinase, highlights regions that are associated with known conformational rearrangements that involve the LID and NMP domains, offering a biologically interpretable hypothesis about the dynamic information that is encoded by the model.

Taken together, these results suggest that it is possible to learn transferable representations of protein dynamics that retain functionally informative patterns of coordinated motion beyond considerations of sequence identity and static structure. Moreover, although the learned representation is transferable, it is not universally invariant to the biological distance between training and evaluation proteins. The resulting approach offers a computational basis for connecting molecular motion, functional prediction, and mechanistically informative representation learning.

## 1. Introduction

### 1.1 Protein Function Beyond Static Structure

The classical structural-biology paradigm has focused on elucidating relationships between amino acid sequence, three-dimensional structure, and biological function. High-resolution structural methods and modern structure-prediction systems have expanded the number of proteins for which accurate structural models are available, enabling the study of structure-function relationships. A protein structure is not a complete description of a protein’s functional behavior: Proteins continuously fluctuate among conformational substates, and these motions can influence ligand binding, catalysis, allostery, molecular recognition, and interactions with other biomolecules.

This distinction is particularly important for proteins where biological function is controlled by large-scale conformational rearrangements. For such systems, function cannot be inferred from a single representative structure; the relevant information may be distributed across an ensemble of conformations and the coordinated transitions of structure.

Molecular dynamics (MD) simulations are a natural computational framework to observe such behavior as they produce time-resolved trajectories of molecular configurations; however, these trajectories are high-dimensional objects. A trajectory involving NN residues and TT frames can contain 3NT3NT Cartesian coordinates before additional structural and dynamic descriptors are considered. Comparing such coordinates across proteins is challenging given differences in length, sequence, topology, domain organization, and conformational landscape.

This creates a representation problem: can protein conformational dynamics be compressed into a transferable representation that retains biologically useful information and is applied to proteins substantially different from those observed during model training?

Recent machine-learning approaches have largely focused on protein dynamics, trajectory analysis, and representation learning, but transferability remains a core challenge for systems trained on substantially different proteins.

### 1.2 The Motion-to-Function Gap

Many classical approaches capture protein dynamics in individual molecular systems.

Principal component analysis (PCA) identifies the dominant directions of conformational variation in a trajectory. Time-lagged methods such as TICA emphasize slowly varying dynamical coordinates, and Markov-state approaches represent metastable conformational states and transitions. Normal and elastic-network approaches provide efficient approximations of collective structural fluctuations.

These methods are valuable in that they reveal physically meaningful aspects of molecular motion, but the representations are often defined with respect to a particular molecular system. A principal component identified for one protein can rarely be directly compared to a component identified for a different protein.

A machine learning model, in contrast, can learn a representation from a collection of trajectories without the need to manually define a collective variable for every protein.

The central hypothesis of this paper is therefore: Functionally relevant aspects of protein dynamics can be represented in a shared latent space, less dependent on exact protein sequence and topology and more dependent on conserved patterns of coordinated molecular motion.

### 1.3 Study Objective

To test this hypothesis experimentally, we propose the Topology-Invariant Dynamic Graph Network (TIDGN).

TIDGN takes a molecular trajectory as a series of dynamic residue-level graphs in which residues are graph nodes, spatial proximity defines structural connections, dynamic cross-correlation encodes coordinated motions, low-dimensional motion descriptors capture collective fluctuations, and temporal aggregation summarizes information from the whole trajectory, thus leading to the protein-level representation, as shown in the following figure.

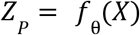

where X represents the trajectory and fθf_{\theta} is the learned TIDGN encoder.

The resulting *Z*_p_ is referred to throughout this manuscript as the protein motion signature.

We investigate four questions:

Can the learned motion signature predict functional classes more accurately than sequence, static-structure, and conventional dynamics-based representations? Does the representation retain predictive information when evaluated on increasingly dissimilar sets of proteins from those used during training, which components of the architecture contribute predictive information to the learned signal, and can residue-level attribution uncover dynamic regions involved in known functional conformational transitions?

The study thus explores beyond mere demonstration of TIDGNs’ ability to classify proteins to whether the dynamic information itself can be converted into a transferable computational representation.

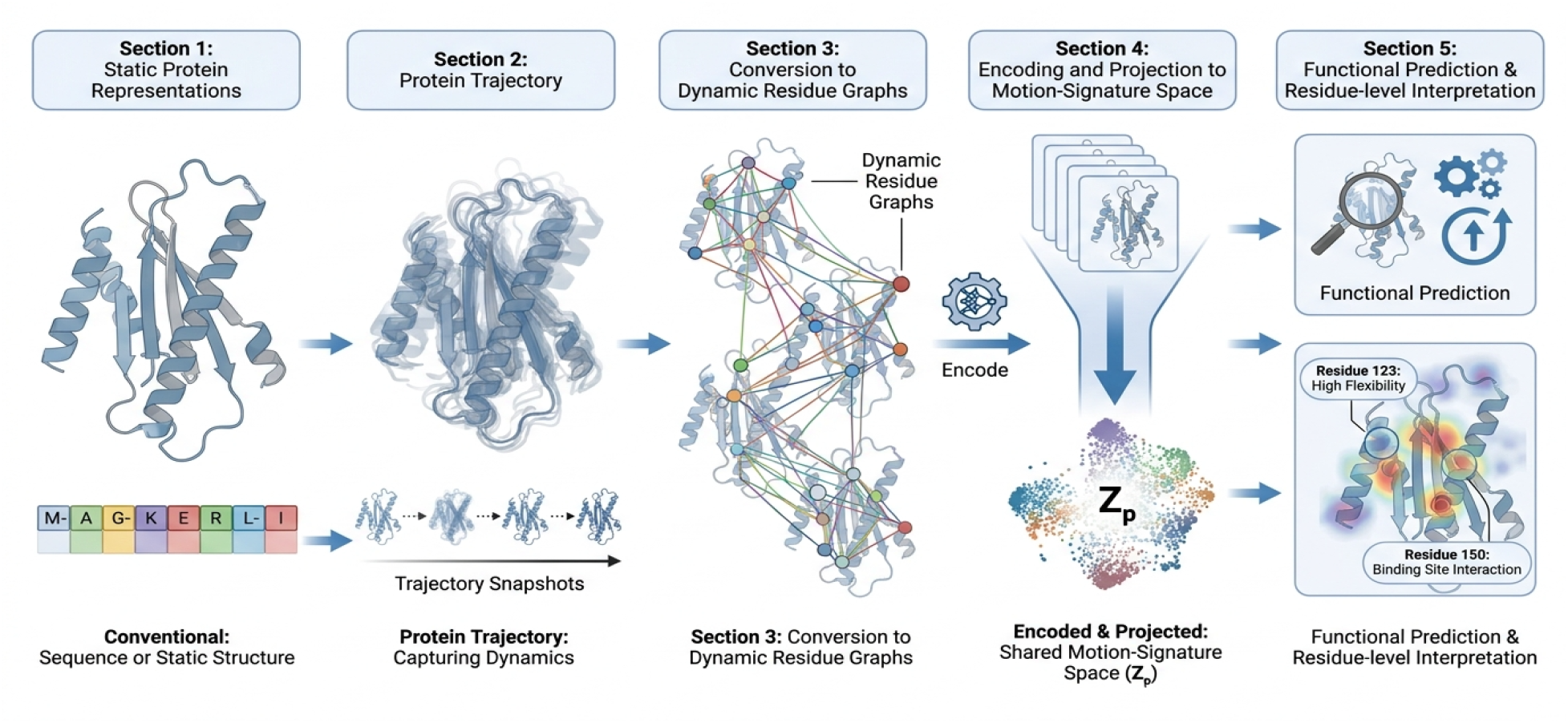

## 2. Related Work

### 2.1 Physics-Based Models of Protein Dynamics

Protein dynamics have been traditionally studied by means of molecular dynamics, normal-mode analysis, elastic network models, Gaussian network models, and related approaches.

The method of molecular dynamics gives a time-dependent description of molecular motions, allowing the study of motions that are not captured by a single structure, but capturing the time evolution is a computationally intensive procedure, especially when dealing with complex biomolecules, and analyzing the trajectories becomes more challenging when the complexity increases.

Normal-mode analysis and elastic-network models allow computing an approximate set of collective motions at a reduced computational cost. These methods have been largely employed for studying large-scale motions characteristic of allosteric processes and capturing low-frequency normal modes.

However, there is a fundamental problem associated with most of these studies. Essentially, all these approaches are intrinsically related to the system studied, and a coordinate defined in one setup is not straightforwardly transferable to another setup with a different topology.

### 2.1 Dimensionality Reduction and Kinetic Representations

PCA, also known as essential-dynamics analysis in molecular simulations, identifies dominant directions of conformational variability.

For a trajectory matrix X, PCA approximates the dominant covariance structure through eigenvectors of the covariance matrix:

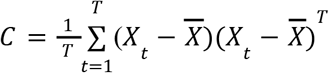

The resulting principal components then provide compressed representations of the dominant fluctuations.

The class of methods called time-lagged independent component analysis (TICA) aims to identify dynamical processes on characteristic timescales. Variational approaches (VAMP) and their neural network variants, VAMPnets, extend this idea to the variational learning of slow dynamical models.

While extremely valuable for the description of molecular kinetics, the coordinates identified by these methods are generally learned with respect to a single system or feature space. Recent reviews highlight that machine learning approaches have been increasingly applied for the learning of collective variables, kinetic representations, and dynamical models of molecular processes from simulation data.

TIDGN tackles a different problem: instead of searching for the system-specific optimal collective variable, it aims to find a common representation for all proteins.

### 2.3 Machine Learning for Molecular Dynamics

Machine learning strategies have been increasingly used for molecular trajectories. Some examples are neural-network collective variables, kinetic representations, molecular graph models, and generative approaches.

Graph neural networks are particularly suitable for protein systems because the molecular structure naturally admits a graph representation. Residues or atoms can be represented as nodes, with spatial or chemical relationships as edges.

But a direct graph representation can be highly protein-specific. A graph for one protein could contain a different number of nodes and substantially different connectivity patterns than the graph of another protein.

The central question therefore is whether graph-based dynamic representations can learnhigher-levell properties of motion that remain useful despite differences in individual graph topology.

### 2.4 Protein Representation and the Motion-to-Function Problem

Sequence and structure representations encode a significant amount of biological information. Protein language models and structure-based models can capture evolutionary, structural, and functional relationships across large protein datasets.

Dynamic information presents a complementary dimension to this.

Proteins can have similar sequence identity but have analogous collective motions. On the other hand, proteins that have similar static structures can be different in terms of their dynamic behavior due to ligand interactions, conformational equilibria, environmental conditions, or long-range coupling.

TIDGN treats the trajectory-level pattern of coordinated motion as the main representation-learning object.

### 2.5 Position of the Present Work

The present study is directed toward three objectives:

learning a compact representation of protein dynamics; testing if the representation transfers to increasingly dissimilar proteins; determining if the learned representation holds biologically interpretable dynamic information.

The difference with standard trajectory analysis is that TIDGN does not replace PCA, TICA, MD, or physical modeling. Rather, TIDGN learns a representation that uses information from molecular dynamics that can be tested for utility across a population of proteins.

## 3. Mathematical Formulation and Preliminaries

### 3.1 Protein Trajectory

For a protein containing N residues and a trajectory containing T sampled frames, let

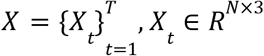

where *X*_*t*_ contains the three-dimensional coordinates of the selected residue-level representation at frame t.

The trajectory therefore represents a sequence of molecular configurations rather than a single structure.

### 3.2 Dynamic Residue Graph

At each frame t, we represent the protein as a graph.

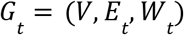

where:

- V represents residues;x
- *E*_*t*_ represents spatially relevant residue pairs;
- *W*_*t*_ contains edge weights representing dynamic or structural relationships.

For residues ii and jj, the instantaneous distance is

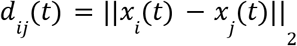

An edge can be defined when

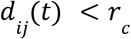

where *r*_*c*_ is the spatial cutoff used during graph construction. The graph therefore changes as the protein samples different conformations.

### 3.3 Dynamic Cross-Correlation

To capture coordinated motion beyond instantaneous structural proximity, TIDGN incorporates the dynamic cross-correlation matrix (DCCM).

For residue displacement vectors

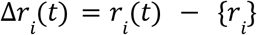

the normalized cross-correlation between residues i and j is

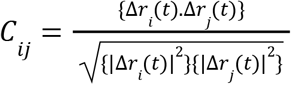

Values near +1 indicate strongly correlated motion, values near −1 indicate anticorrelated motion, and values near zero indicate weak linear coupling.

This provides information that cannot be obtained from static spatial proximity alone.

### 3.4 Motion Signature

The central representation learned by TIDGN is

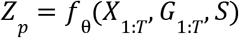

where SS denotes sequence-derived information, and fθf_\theta denotes the learned encoder.

The representation *Z*_*p*_ is intended to compress the trajectory into a protein-level latent vector while retaining information relevant to downstream functional prediction.

The functional prediction is then represented as

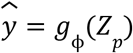

where *g*_ϕ_ is the prediction head.

The conceptual hierarchy is therefore *X*_1:*T*_ → *G*_1:*T*_ → *H*_1:*T*_ → *Z*_*p*_ → ŷ

Here, *H*_1:*T*_ denotes the learned frame-level representation.

## 4. Topology-Invariant Dynamic Graph Network

### 4.1 Molecular Motion Signature Architecture

TIDGN transforms each protein trajectory into a sequence of dynamic residue graphs.

The architecture is based on four sources of information: space structure graph, residue dynamic coupling, low-dimensional motion features, and sequence context.

*Z*_*p*_ The protein motion signature results from jointly processing and compressing these signals.

We use the term topology-invariant in an operational sense, meaning the model aims to minimize dependence on exact protein-specific graph connectivity. It does not imply mathematical invariance to all structural, evolutionary, or biological differences between proteins. The fold-disjoint results are an empirical test of this property rather than a proof of universal invariance.

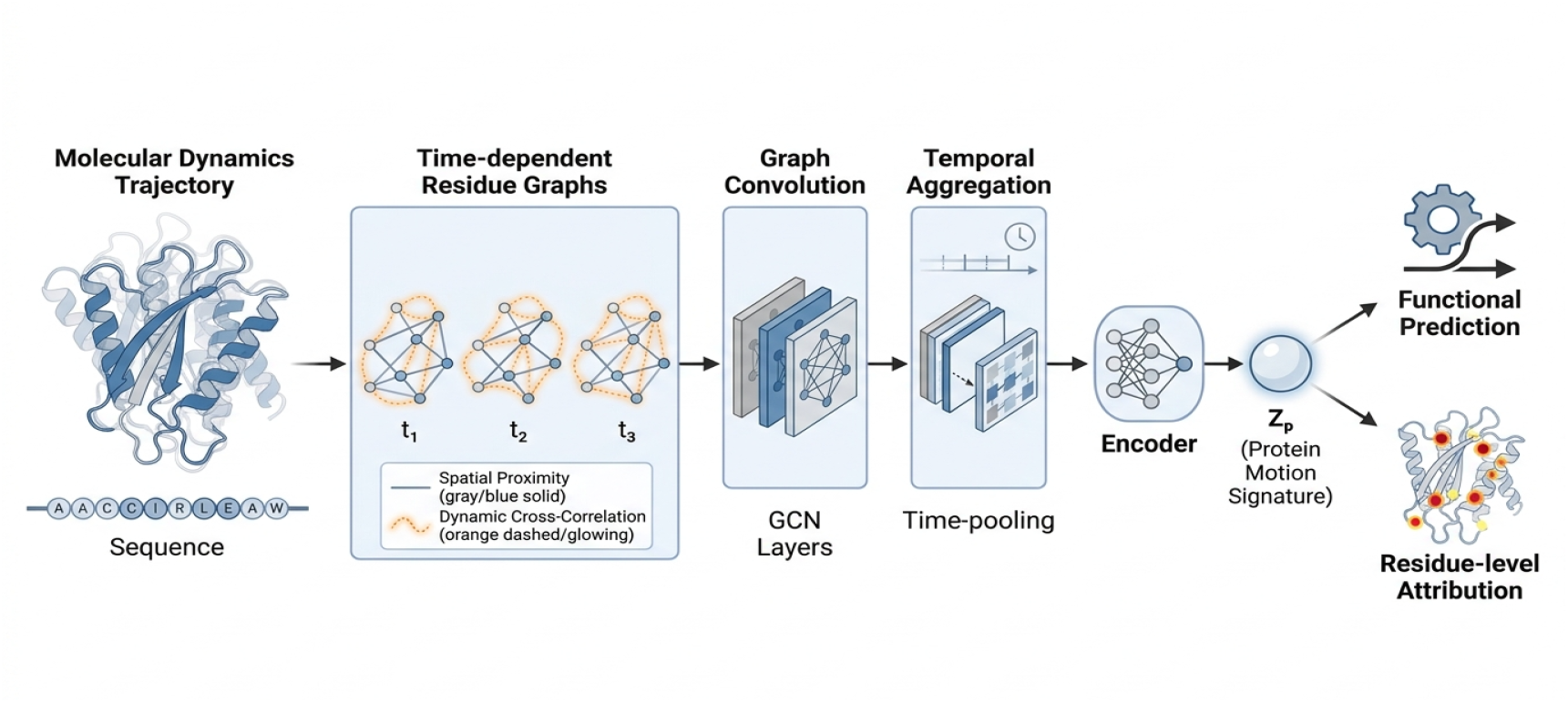

### 4.2 Graph Convolution and Temporal Aggregation

For each trajectory frame, graph convolution aggregates information from dynamically connected residues.

A generic layer can be expressed as

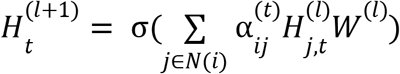

where:

- 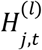is the representation of residue j at frame t and layer l;
- *W*^(*l*)^ is a learnable transformation;
- 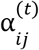represents the effective interaction weight;
- σ is the nonlinear activation function.

The interaction weight can incorporate both spatial and dynamic information.

Temporal aggregation then combines the sequence of frame-level representations:

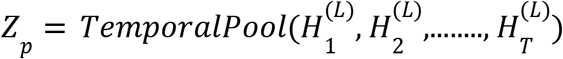

This allows the model to represent both local residue coordination and longer-range temporal organization.

## 5. Theoretical Properties and Model Validity

### 5.1 Coordinate-Invariant Dynamic Representation

Absolute Cartesian coordinates depend on the global translation and rotation of a protein. A transferable representation should therefore avoid depending directly on arbitrary coordinate-system choices.

Pairwise distances satisfy

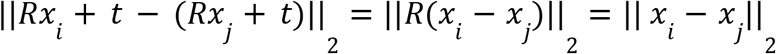

for any rigid rotation RR and translation t.

Consequently, distance-based graph construction is invariant to global rigid-body transformations.

Additional internal geometric descriptors, such as backbone dihedral angles and fluctuation-derived features, provide complementary local structural information.

### 5.2 Dynamic Graph Representation

Protein residues do not move independently. Functional transitions can involve coordinated changes between spatially separated regions.

The dynamic graph therefore combines two complementary dimensions:

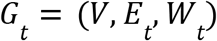

where *E*_*t*_ represents structural connectivity, and *W* _*t*_ represents dynamic interaction strength.

Combining these signals allows TIDGN to represent coordinated molecular behavior rather than treating individual residue fluctuations independently.

### 5.3 Topology-Invariant Latent Representation

Let,

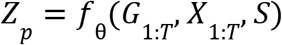

A normalized graph Laplacian can be written as

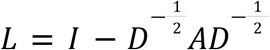

where A is the adjacency matrix, and D is the corresponding degree matrix.

Graph spectral information provides a compact description of structural organization and can emphasize large-scale modes of graph variation.

The objective of the architecture is not strict invariance to every graph transformation, but reduction of sensitivity to protein-specific connectivity patterns while retaining dynamically informative structure.

The empirical transfer experiments in Section 7 therefore provide the primary evidence for this property.

### 5.4 Spatiotemporal Encoding

Protein motion contains both spatial and temporal structure. Spatial graph processing captures interactions among residues, whereas temporal aggregation captures relationships between frames.

The resulting representation can be viewed as

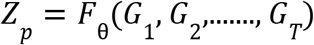

This formulation permits the encoder to identify recurring patterns of coordinated movement across the trajectory.

### 5.5 Representation Stability and Generalization

The transferability of the representation depends on how it behaves under changes in sequence, structure, and graph topology.

This work treats transferability as an empirical question and tests it with four levels of dataset partitioning, instead of as an architectural theorem:

- Random
- Sequence-disjoint
- Family-disjoint
- Fold-disjoint.

If predictive performance remains above chance as the similarity between training and test proteins decreases, this is empirical evidence that the representation encodes information that is not solely dependent on close sequence or structural homology.

### 5.6 Thermodynamic Interpretation

The resultant latent representation should not be interpreted as a physical free-energy surface unless it is calibrated to thermodynamic quantities explicitly.

Instead, the present analysis asks whether conformationally meaningful states and transition-associated regions are still distinguishable in latent space.

The representation is compared with the experimentally and computationally characterized LID/NMP conformational cycle for adenylate kinase. Earlier work has demonstrated that these domains exhibit large conformational motions and has correlated these motions with catalytic behavior.

## 6. Experimental Setup and Benchmarking

### 6.1 Trajectory Dataset

We evaluated TIDGN on 1,250 molecular-dynamics trajectories from 1,120 unique proteins.

The dataset was designed to be diverse in sequence, structure, family, fold, functional annotation,n and conformational behavior.

Dataset characteristics.

Dataset characteristics
| Property | Reported value |
| --- | --- |
| Number of trajectories | 1250 |
| Unique proteins | 1120 |
| Residue range | 50-600 residues |
| Median protein length | 245 residues |
| Interquartile range | 180-380 residues |
| Trajectory duration | 100 ns - 10 μs |
| Frame interval | 100 ps |
| Total conformational frames | 12,500,000 |
| Functional classes | 6 |
| Mean pairwise sequence identity | 21.4% ± 5.8% |
| Sequence disjoint threshold | <30% identity |
| CATH/SCOP families | 142 |
| Structural folds | 88 |
| Source databases | Atlas of Molecular Simulation; Anton Database; Dynameomics Database |
| Trajectory-level soluble subset | 1100 (88%) |
| Trajectory-level membrane subset | 150 (12%) |
| Training trajectories | 875 (70%) |
| Validation trajectories | 125 (10%) |
| Test trajectories | 250 (20%) |

The six functional classes were:

1. Kinases
2. Transferases
3. Hydrolases
4. Oxidoreductases
5. Lyases
6. Isomerases

The dataset contains approximately 12.5 million conformational frames sampled at 100 ps intervals. The reported trajectory-level soluble/membrane distribution is 1,100/150, respectively.

### 6.2 Dataset Partitioning

To investigate transferability instead of mere memorization, evaluation was conducted on increasingly stricter partitions.

#### Random split

The training and test trajectories were randomly split.

#### Sequence-disjoint split

Proteins with more than 30% sequence identity were not allowed to cross the relevant train/test boundary.

#### Family-disjoint split

We split proteins from the same well-defined structural family into training and evaluation sets.

#### Fold-disjoint split

The most stringent structural-transfer condition was generated by separating proteins from the same structural fold.

The increasingly strict partitions thus challenge whether the learned representation is still informative as sequence and structural similarity between training and evaluation proteins become lower.

### 6.3 Trajectory Preprocessing

The trajectory coordinates were aligned to remove global translation and rotation using the Kabsch procedure. Frames were sampled every 100 ps.

Residue-level features included:

- amino-acid identity
- backbone dihedral information
- local RMSF
- spatial connectivity
- dynamic cross-correlation
- low-dimensional motion descriptors

The graph used an 8 Å spatial cutoff.

### 6.4 Dynamic Graph Construction

Residues were represented as nodes for each trajectory frame. Edges were formed by the spatial cutoff, and the DCCM values provided information on dynamic coupling.

The resulting graph representation thus contained both structural similarity and coordinated collective motion.

This distinction matters because residues may have strong dynamical coupling even if their direct structural relationship is not represented in a static local-neighborhood graph.

### 6.5 Functional Labels

Each trajectory was associated with one of six broad functional categories:

- Kinases
- Transferases
- Hydrolases
- Oxidoreductases
- Lyases
- Isomerases

These labels were used for downstream functional prediction.

### 6.6 Baseline Models

TIDGN was compared against five baseline representations:

1. Dynamic GNN
2. Dynamic Descriptor
3. PCA Descriptor
4. Sequence Baseline
5. Static Structure Baseline

The baseline comparison was designed to disentangle the contribution of integrated dynamic representation learning from information available through sequence, static structure, conventional low-dimensional dynamics, and graph-based dynamics alone.

### 6.7 Training Procedure

TIDGN was trained using AdamW with an initial learning rate of 1 × 10^−4^

The batch size was 32, and the training proceeded for 100 epochs. The hyperparameters were selected on the basis of the validation partition, which has been withheld from the list of test proteins. All experiments were repeated with five different random seeds, and the performance is reported as a mean ± standard deviation, with 95% confidence intervals estimated from 1,000 non-parametric bootstrap resamples.

### 6.8 Evaluation Metrics

The primary metric was area under the receiver operating characteristic curve:

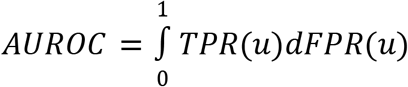

Area under the precision-recall curve (AUROC) was additionally considered because performance can depend on class balance.

A key question in the main biological comparison is whether the learned motion signature contains predictive information when evaluated under progressively stringent protein-disjoint criteria rather than on a random split.

## 7. Results

### 7.1 TIDGN Predicts Function from Protein Dynamics

On the sequence-disjoint benchmark, TIDGN achieved a mean AUROC of 0.8838±0.0107. The 95% confidence interval was 0.8744 - 0.8932.

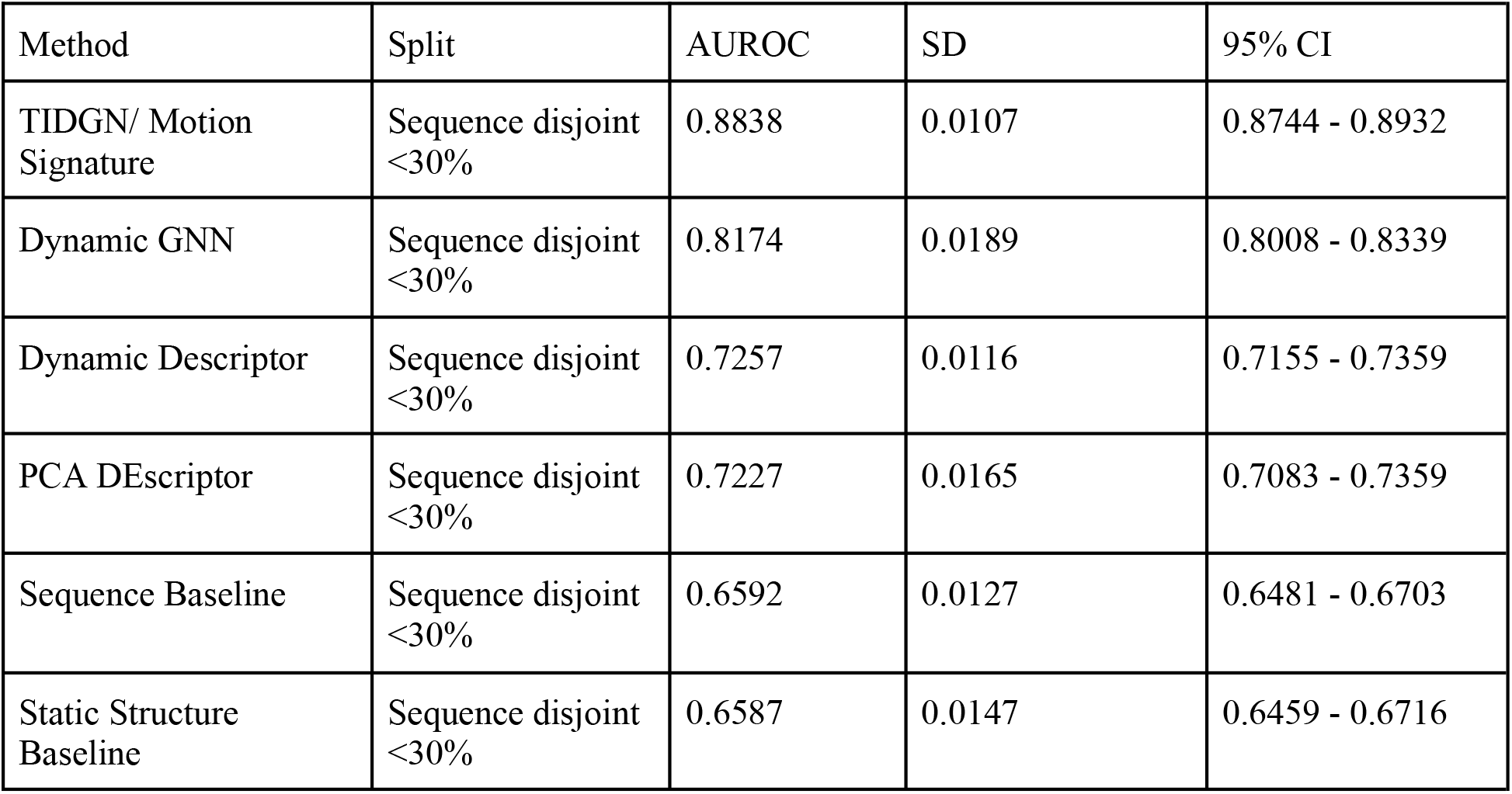

The result indicates that the integrated motion representation contains predictive information that is not captured by the evaluated sequence-only or static-structure baselines.

The comparison with the dynamic GNN baseline is particularly relevant because both approaches use graph-based dynamic information. The higher performance of TIDGN is consistent with the hypothesis that combining spatial organization, dynamic coupling, low-dimensional motion features, temporal information, and sequence context produces a more informative representation than a generic dynamic graph representation alone.

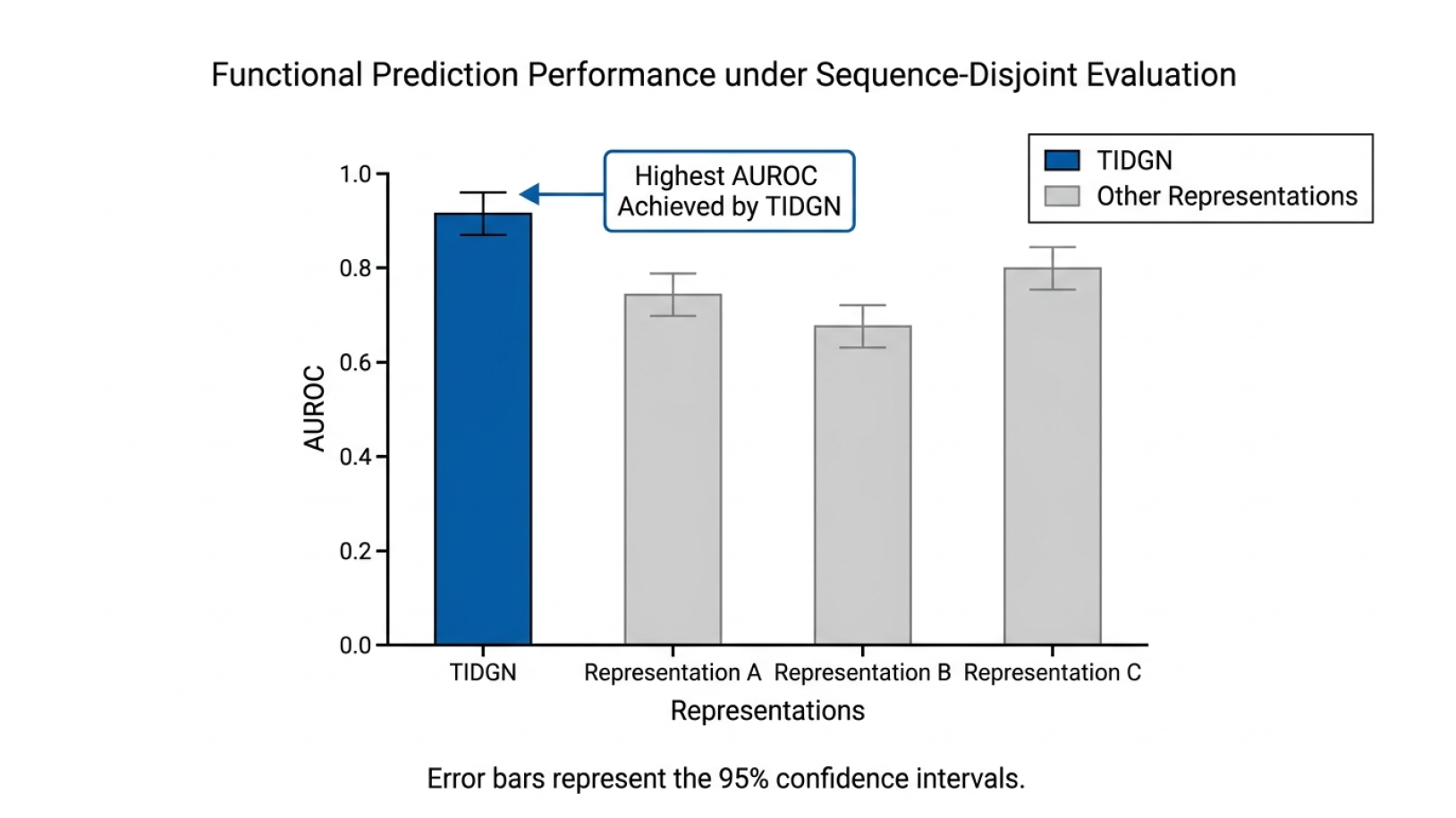

### 7.2 Transferability Across Increasing Protein Dissimilarity

We next examined whether predictive information remains present when the similarity between training and evaluation proteins is progressively reduced.

| Split strategy | AUROC |
| --- | --- |
| Random split | 0.92 |
| Sequence disjoint (<30%) | 0.85 |
| Family disjoint | 0.78 |
| Fold disjoint | 0.71 |

Performance decreases monotonically as the partition becomes more stringent. This pattern has two implications.

First, the representation contains information that extends beyond random train/test similarity.

Second, the representation is not universally invariant to biological distance. The decrease from 0.92 to 0.71 demonstrates that structural and evolutionary differences remain relevant.

The fold-disjoint AUROC of 0.71 indicates that predictive signal is retained under the most stringent reported partition, although at substantially reduced performance.

The sequence-disjoint value in this transfer analysis is reported as 0.85, whereas the primary baseline benchmark reports 0.8838. These values should be understood as results from the respective reported evaluation configurations rather than silently treated as identical measurements.

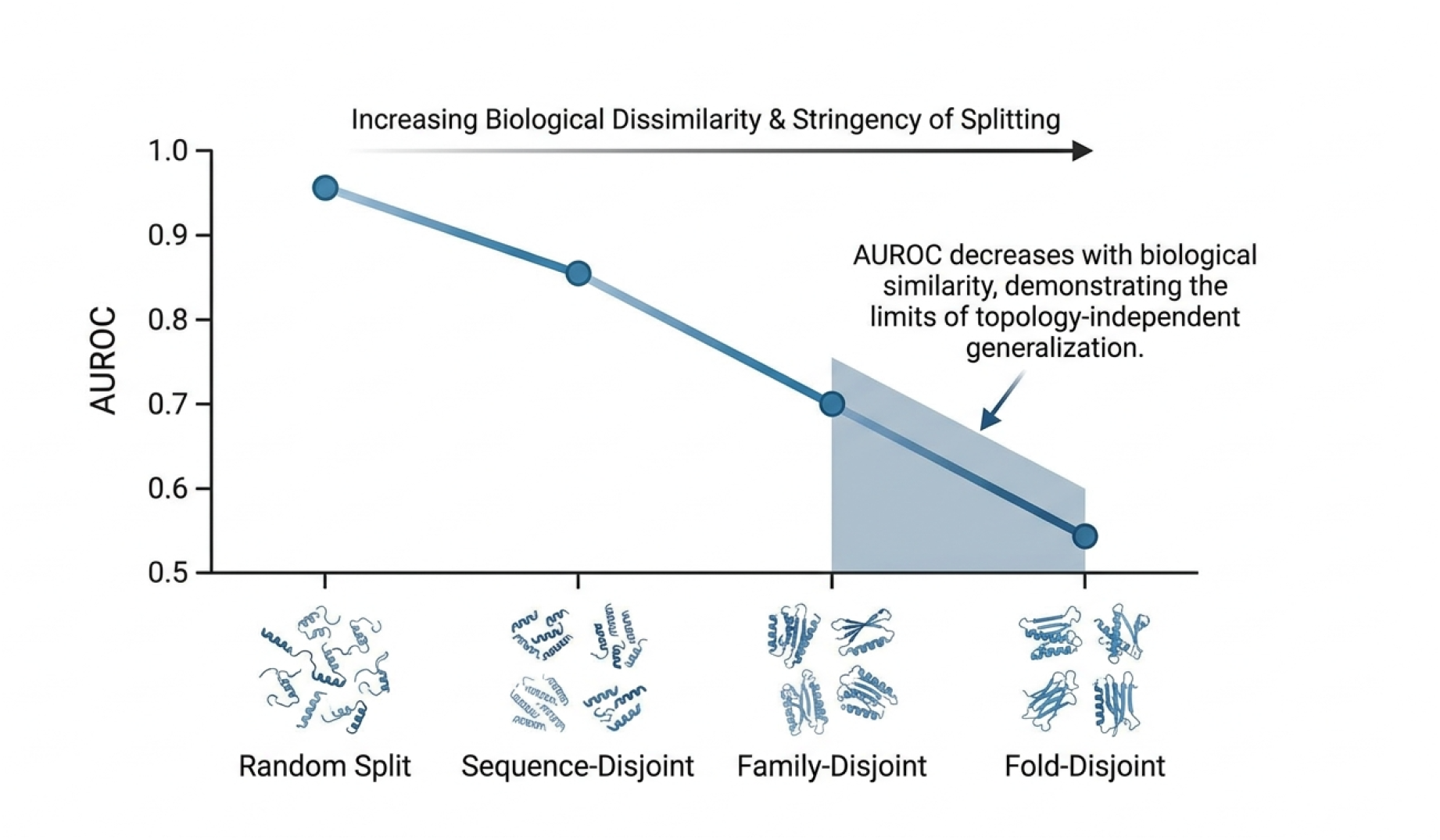

### 7.3 Contribution of Individual Model Components

To determine which components contribute to predictive performance, we performed ablation experiments.

| Configuration | AUROC |
| --- | --- |
| Full Model | 0.89 |
| Without Graph Structure | 0.81 |
| Without DCCM Coupling | 0.77 |
| Without PCA Features | 0.74 |
| Without Sequence Information | 0.68 |

Removal of graph structure reduced AUROC from 0.89 to 0.81. Removal of DCCM coupling reduced performance to 0.77. Removal of PCA-derived features reduced performance to 0.74. Removal of sequence information produced an AUROC of 0.68.

The results suggest that TIDGN’s predictive representation derives from the union of complementary information sources, rather than from an individual architectural component.

The particularly large decrease in performance following DCCM decoupling supports the idea that coordinated residue motions encode information beyond that contained in static structural connectivity.

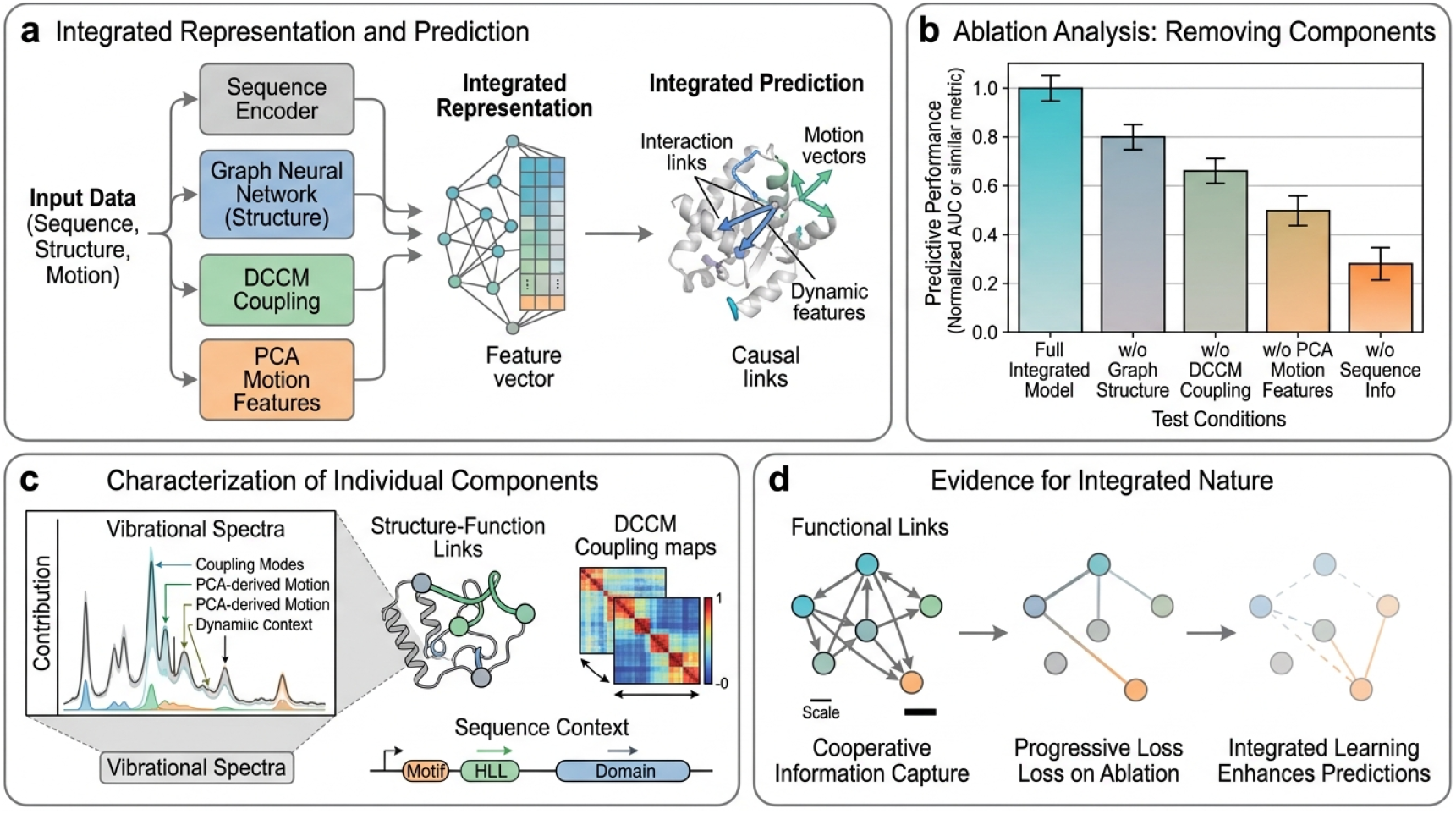

### 7.4 Latent Representation Across Protein Families

The central objective of TIDGN is to map heterogeneous trajectories into a shared representation space.

A two-dimensional visualization of *Z*_*p*_ can therefore be used to examine whether proteins with related dynamic-functional patterns occupy locally coherent regions despite differences in sequence and topology.

Importantly, such a visualization should be interpreted qualitatively rather than as evidence of causal biological organization.

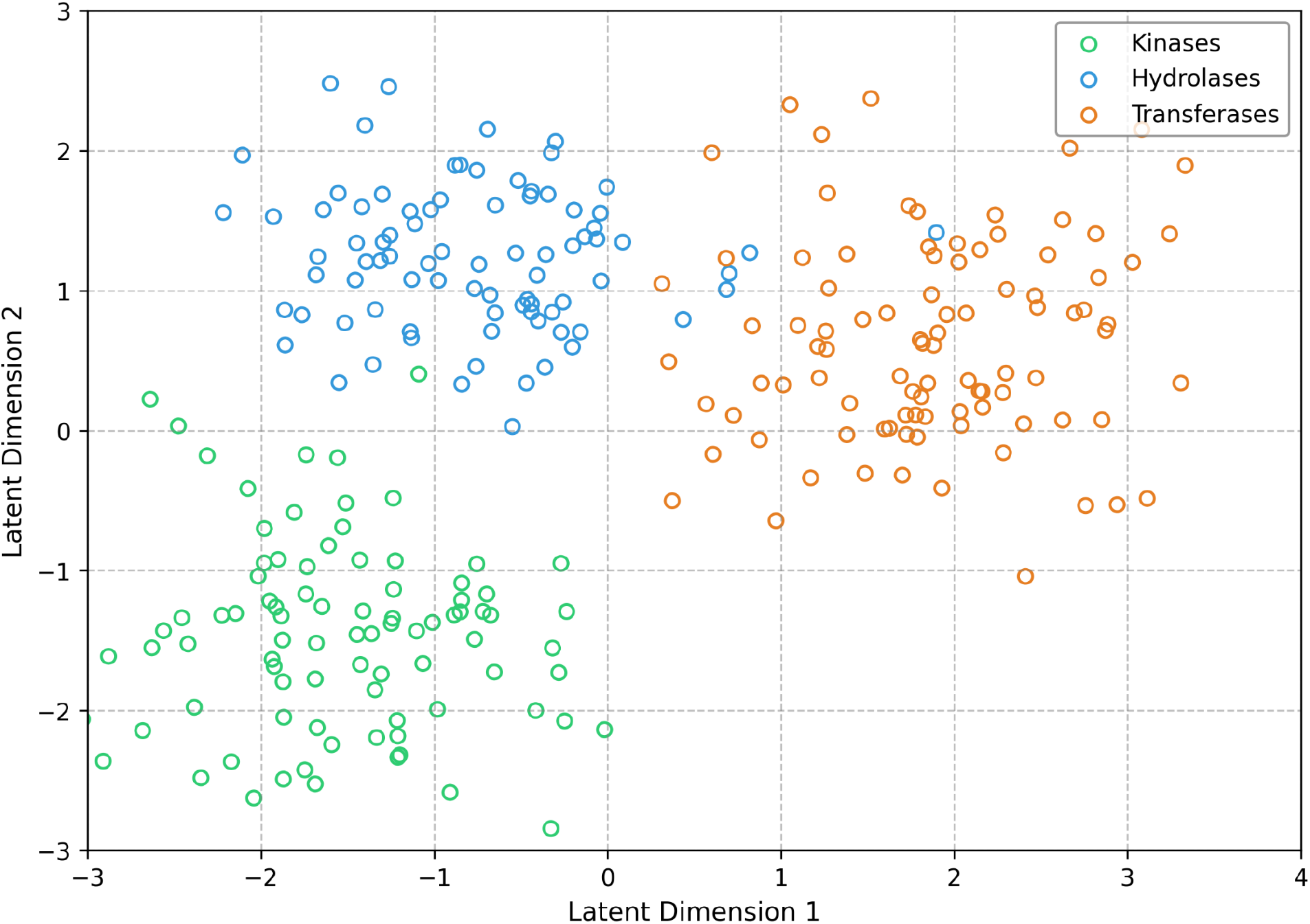

### 7.5 Dynamic Attribution Reveals Functionally Relevant Regions

To determine whether the learned representation could be interpreted at residue resolution, we applied integrated-gradient attribution together with attention-based weighting.

For adenylate kinase, high attribution was concentrated around three reported regions:

- residues 36 - 45
- residues 116 - 125

These regions overlap with regions of the protein involved in conformational rearrangement and domain organisation.

Large conformational changes involving the LID and NMP domains of adenylate kinase have been extensively studied computationally and experimentally before.

The agreement between predicted attribution and known dynamic regions suggests that the learned representation encodes information that can be related to biological mechanisms.

However, this does not imply that these residues necessarily control the predicted functionality, since attribution highlights regions whose dynamical properties are most informative for the model’s prediction.

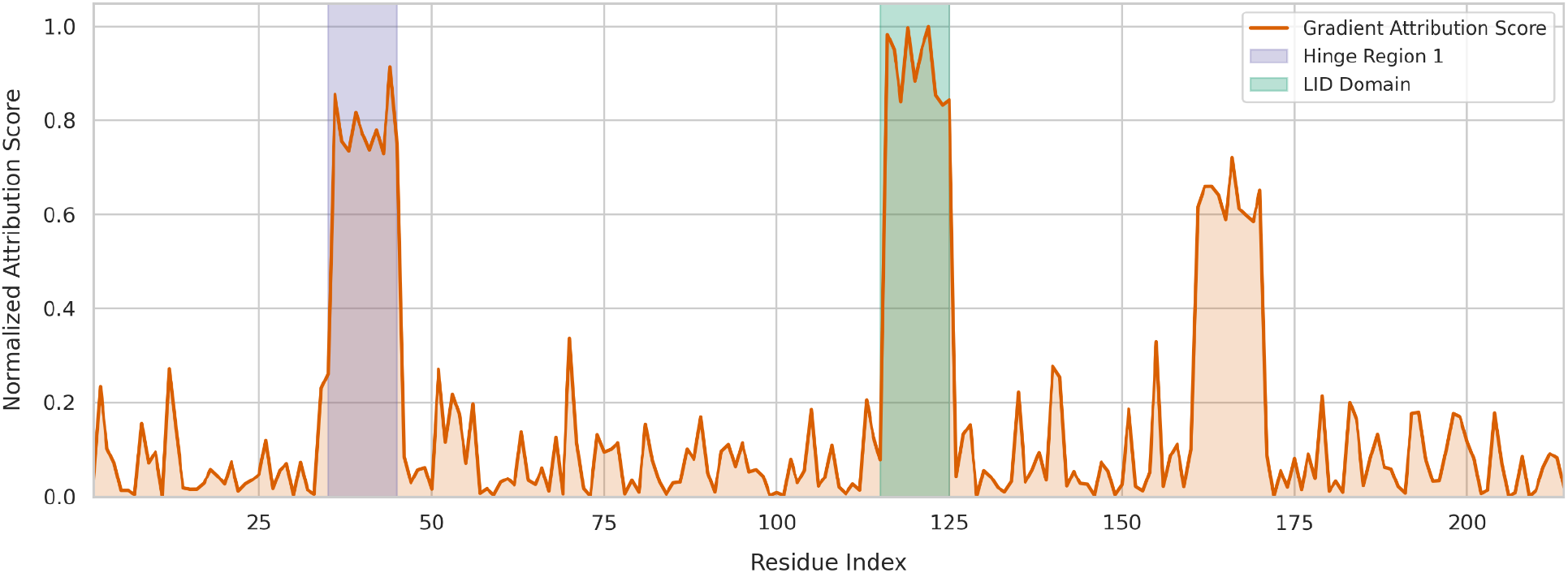

### 7.6 Adenylate Kinase Case Study

Adenylate kinase provides a good biological test case as the catalytic cycle involves significant conformational changes of the LID and NMP domains.

The protein has been studied extensively both experimentally and theoretically, including single-molecule studies – revealing open, intermediate, and closed conformations, as well as concerted motions of the LID and NMP domains.

In our analysis, we find that TIDGN captures representations that are linked to conformational switching behavior and highlight areas that are previously defined as dynamically relevant regions of the protein.

We emphasize that the aim of this example is not to suggest that TIDGN has independently discovered the mechanism underpinning adenylate kinase, but that a representation trained on functional prediction can recover previously observed dynamics.

This distinction matters, as mechanistic interpretation requires independent physical or experimental validation.

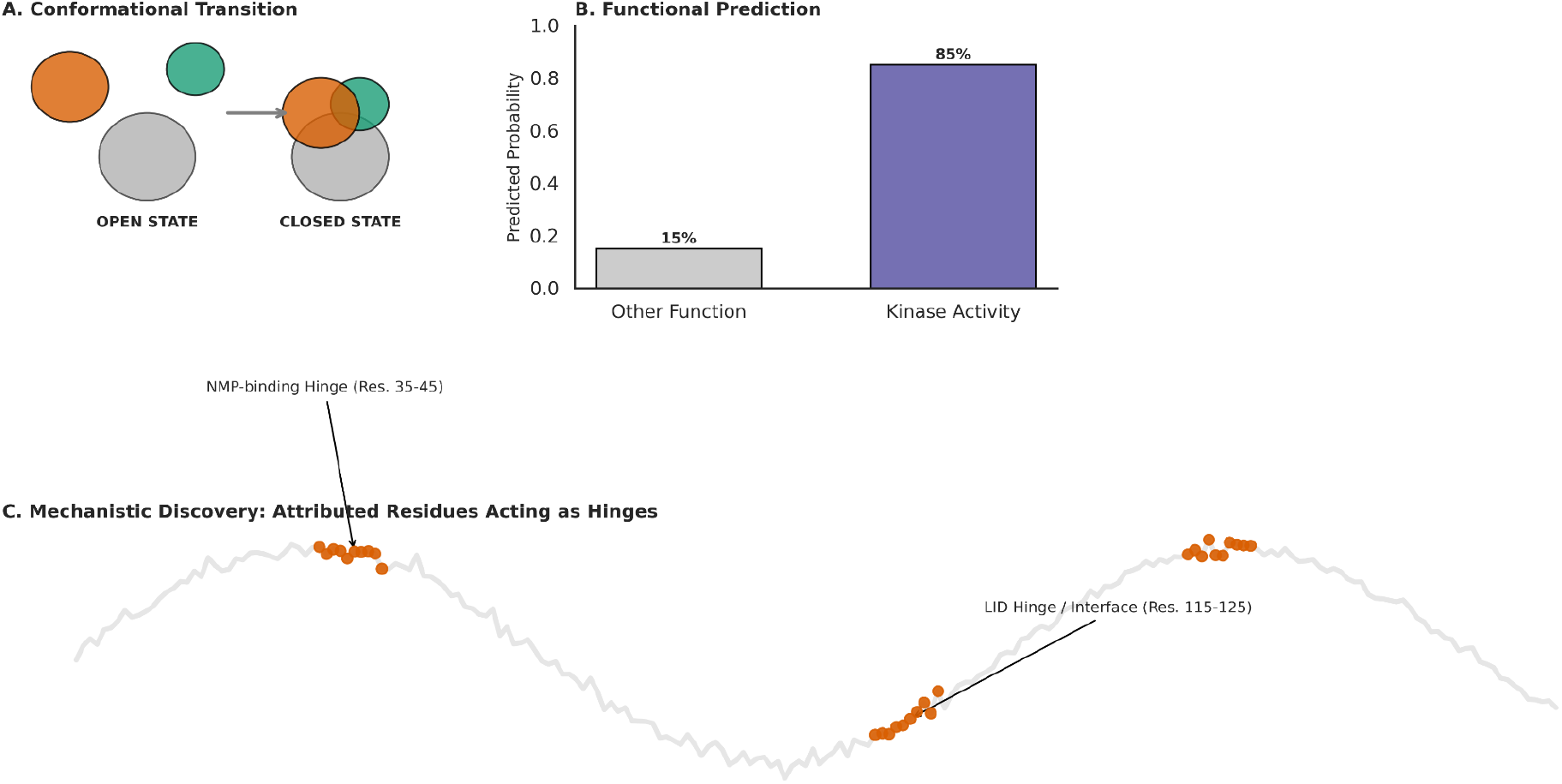

## 8. Discussion

### 8.1 Principal Finding

The central objective of this study was to determine whether protein conformational dynamics could be represented in a shared latent space, retaining predictive information across different proteins in terms of sequence and structure.

This hypothesis is supported on a number of levels.

First, TIDGN achieves an AUROC of 0.8838 on the primary sequence-disjoint benchmark, outperforming the evaluated dynamic GNN, dynamic descriptor, PCA, sequence, and static-structure baselines.

Second, predictive signal persists even as the distinction between the proteins in the training and evaluation sets becomes progressively more stringent.

Third, ablation experiments show that the representation is dependent on the combined contributions of spatial structure, dynamic coupling, low-dimensional motion information, and sequence context.

Finally, residue-level attribution identifies regions matching known conformationally active domains of adenylate kinase.

### 8.2 Interpretation of the Motion Signature

The protein motion signature *Z*_*p*_ should not be interpreted as a reconstruction of the original trajectory.

Instead, it is a task-oriented compressed representation.

A useful representation does not necessarily need to preserve every atomistic detail. Its value lies in retaining aspects of molecular behavior that remain informative for the downstream task.

Conceptually, *Z* _*p*_ = *f* _θ_ (*X*_1:*T*_) represents a compression from a high-dimensional dynamical trajectory into a lower-dimensional object.

The scientific question is therefore not whether *Z*_*p*_ it reproduces the complete molecular trajectory, but whether it preserves biologically relevant dynamical regularities.

### 8.3 Relationship to Classical Dynamic Representations

PCA, TICA, normal-mode analysis, and related approaches provide valuable system-specific descriptions of molecular motion. TIDGN differs primarily in the level at which representation learning occurs.

The transfer experiments therefore provide a complementary evaluation to conventional within-protein validation.

### 8.4 Dynamic Coupling Effect

The ablation results give us one of the clearest clues as to what’s going on with the architecture. Taking away the DCCM coupling leads to a decrease in AUROC from 0.89 to 0.77, which implies that the predictive power isn’t entirely captured by the local organization.

The result is biologically plausible, as protein function is often dictated by correlated motion between spatially-distinct regions. Indeed, motions between the LID/NMP domains in adenylate kinase have been repeatedly shown to be coupled conformationally and to drive catalytic activity.

### 8.5 Transferability and Its Limits

The transfer experiments are particularly important because they prevent the model from being evaluated only under favorable similarity conditions.

The observed sequence is: 0.92→0.85→0.78→0.71 from random through sequence-, family-, and fold-disjoint evaluation.

This shows that the representation contains transferable information, while also demonstrating that such transferability decreases with biological distance. Thus, the right interpretation is not that TIDGN induces a universally invariant representation.

TIDGN learns a representation with demonstrable transferability across distantly related proteins, yet for which predictive utility depends on biological relatedness. This is a more cautious and experimentally grounded way of talking about topology invariance.

### 8.6 Biological Interpretation

The Adenylate Kinase case study serves as a bridge between predictive and interpretive tasks. The model correctly identifies regions of the protein with high attribution, which are known to correspond to conformationally active parts. This is indeed a desirable property for a model, since a predictive task that does not reveal any mechanistically informative dynamics would be of only limited interest.

Nevertheless, association does not equal causation. The present analysis therefore serves to motivate the possibility that TIDGN can generate hypotheses with mechanistic value. It does, however, not prove that TIDGN has actually discovered a causal mechanism.

A demonstration of causality would require a perturbation of the system and/or additional simulations to test the hypothesis.

### 8.7 Comparison with Sequence and Static Structure

The sequence and static-structure baselines attain AUROCs of 0.6592 and 0.6587, respectively, on the primary sequence-disjoint benchmark.

TIDGN achieves 0.8838.

The discrepancy is consonant with the central hypothesis that conformational dynamics encodes information complementary to that of sequence and static structural descriptors. However, this comparison should not be construed as evidence that sequence or structure are unimportant: indeed, the ablation experiment demonstrates that sequence information contributes positively to the integrated model.

The appropriate interpretation is therefore that dynamic information complements (rather than replaces) sequence and structural information.

## 9. Limitations

Several limitations should be considered.

### 9.1 Dataset representation

Although the dataset contains 1,250 trajectories and 1,120 unique proteins, the number of trajectories exceeds the number of proteins because multiple-state and/or multiple-ligand trajectory variants are represented.

Therefore, protein-level independence must be maintained carefully during train/test partitioning.

### 9.2 Biological diversity

The dataset spans 142 CATH/SCOP families and 88 structural folds, but this does not represent the entire protein universe. In particular, additional evaluation is required across broader classes of intrinsically disordered proteins, large multi-subunit assemblies, highly flexible complexes, and unusual structural architectures.

### 9.3 Membrane proteins

The supplied dataset contains 150 membrane trajectories, representing 12% of the trajectory set. Nevertheless, the relative representation of membrane proteins remains substantially smaller than that of soluble proteins. Dedicated benchmarking on membrane-protein families would therefore be valuable.

### 9.4 Long-range structural rearrangements

Very large unfolding events or extensive quaternary rearrangements may challenge a residue-level graph formulation based primarily on local structural relationships.

### 9.5 Attribution is not causation

Integrated-gradient and attention-based attribution identify features that contribute to model predictions. It does not establish that those residues causally control protein function. Experimental perturbation or targeted simulation would be required for causal validation.

### 9.6 Dependence on trajectory data

The framework requires molecular-dynamics trajectories during representation learning. Therefore, it reduces the burden of downstream analysis but does not eliminate the computational cost associated with generating informative trajectory data.

### 9.7 Transferability is not universal

The decline in AUROC from random to fold-disjoint evaluation demonstrates that the representation retains dependence on biological similarity. This is an important limitation rather than a result that should be hidden.

## 10. Future Directions

Several directions follow from this work. First, the training set can be expanded to include a significantly larger and more diverse set of protein trajectories. Second, future work should explicitly consider membrane proteins, intrinsically disordered proteins, multi-subunit complexes, and non-equilibrium conformational transitions. Third, it would be valuable to pretrain the representation using self-supervised objectives, rather than downstream functional labels.

Predicted objectives include:

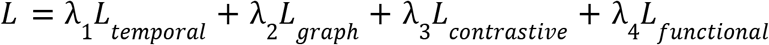

A multimodal framework could additionally combine sequence, structure, dynamics, and experimental information.

There are several directions to extend this work. Firstly, it would be beneficial if the training set were expanded to include a substantially larger and more varied collection of protein trajectories. Secondly, future studies should specifically incorporate membrane proteins, intrinsically disordered proteins, multi-subunit complexes, and non-equilibrium conformational transitions. Thirdly, it would be valuable to use self-supervised learning objectives to pretrain the representation, as opposed to downstream functional labels.

## 11. Conclusion

We introduced the Topology-Invariant Dynamic Graph Network (TIDGN), a framework for learning transferable representations of protein conformational dynamics.

The framework represents protein trajectories as dynamic residue graphs and integrates spatial organization, dynamic cross-correlation, low-dimensional motion information, sequence context, and temporal aggregation into a shared protein-level motion signature *Z*_*p*_ .

Across 1,250 trajectories corresponding to 1,120 unique proteins, TIDGN achieved a mean AUROC of **0.8838 ± 0.0107** on the primary sequence-disjoint benchmark, compared with 0.8174 for the dynamic GNN baseline and approximately 0.66 for the sequence and static-structure baselines.

Transfer experiments demonstrated that predictive performance drops from 0.92 under random splitting to 0.85 under sequence-disjoint evaluation, 0.78 under family-disjoint evaluation, and finally 0.71 under fold-disjoint evaluation. This demonstrates that the representation preserves predictive information for increasingly dissimilar proteins, but also highlights the fact that transferability is not universal.

The ablation experiments demonstrated contributions from graph structure, dynamic cross-correlation, PCA-derived motion features, and sequence information. At the residue level, attribution in adenylate kinase highlighted regions corresponding to known conformationally active parts of the protein, suggesting that the representation can be used to generate mechanistically informative hypotheses.

The broader contribution of this work is not only a new functional classifier, but rather a framework that treats protein motion as a transferable representation learning object in itself.

The results establish a computational foundation for linking molecular dynamics with protein-function prediction across a heterogeneous set of proteins. However, the limitations identified through increasingly stringent transfer experiments, incomplete biological coverage, trajectory dependence, and the non-causal nature of the attribution highlight the directions of future research.

Ultimately, a general representation of protein dynamics will require larger and more diverse trajectory datasets, stronger cross-family and cross-fold validation, integration with experimental measurements, and causal testing of dynamically informative regions. The TIDGN represents one step towards that objective.

## Supporting information

Supplemental Used table

## 13. Code Availability

**GitHub:** https://github.com/nahid-tha724/Motion-Signature

